# Chromosome-level quality scaffolding of brown algal genomes using InstaGRAAL, a proximity ligation-based scaffolder

**DOI:** 10.1101/2019.12.22.882084

**Authors:** Lyam Baudry, Martial Marbouty, Hervé Marie-Nelly, Alexandre Cormier, Nadège Guiglielmoni, Komlan Avia, Yann Loe Mie, Olivier Godfroy, Lieven Sterck, J. Mark Cock, Christophe Zimmer, Susana M. Coelho, Romain Koszul

## Abstract

Hi-C has become a popular technique in recent genome assembly projects. Hi-C exploits contact frequencies between pairs of loci to bridge and order contigs in draft genomes, resulting in chromosome-level assemblies. However, application of this approach is currently hampered by a lack of robust programs that are capable of effectively treating this type of data, particularly open source programs. We developed instaGRAAL, a complete overhaul of the GRAAL program, which has adapted the latter to allow efficient assembly of large genomes. Both GRAAL, and instaGRAAL use a Markov Chain Monte Carlo algorithm to perform Hi-C scaffolding, but instaGRAAL features a number of improvements including a modular polishing approach that optionally integrates independent data. To validate the program, we used it to generate chromosome-level assemblies for two brown algae, *Desmarestia herbacea* and the model *Ectocarpus* sp., and quantified improvements compared to the initial draft for the latter. Overall, instaGRAAL is a program able to generate, using default parameters with minimal human intervention, near-complete assemblies.

## Background

Despite continuous and impressive developments in DNA sequencing technologies, technical challenges remain regarding the assembly of sequence data into full length chromosome assemblies, especially for large genomes [1,2]. Conventional assembly programs and pipelines often encounter difficulty closing gaps in draft genome assemblies caused by regions enriched in repeated elements. At the chromosome level, these programs often incorrectly orient DNA sequences or predict incorrect numbers of chromosomes [3]. Conventional assemblers efficiently generate overlapping set of reads (i.e. contiguous sequences, or contigs) but encounter difficulties linking these contigs together into scaffolds. Consequently, many available genomes feature gaps which need to be bridged to reach a chromosome-level structure. These computational limitations are being addressed thanks to active support from the community and competitions such as GAGE [4] or the Assemblathon [5] but there is as yet no systematic, reliable way of producing near-perfect genome assemblies of guaranteed optimal best quality without a considerable amount of empiric parameter adjustment and manual post-processing evaluation and correction [6].

Recent sequencing projects have typically relied on a combination of independently obtained data such as optical mapping, long read sequencing, and chromosomal conformation capture (3C, Hi-C) to obtain large genome assemblies of high accuracy. The latter procedure derives from techniques aiming at recovering snapshots of the higher-order organization of a genome [7,8]. When applied to genomics, Hi-C-based methods are sometimes referred to as proximity ligation approaches, as they quantify and exploit physical contacts between pairs of DNA segments in a genome to assess their collinearity along a chromosome, and the distance between the segments [9]. Early studies using control datasets demonstrated that Hi-C scaffolds large eukaryotic DNA regions [10–12]. The Hi-C scaffolder GRAAL (Genome Re-Assembly Assessing Likelihood from 3D), a probabilistic tool that uses a Markov Chain Monte Carlo (MCMC) approach was able to generate the first chromosome-level assembly of an incomplete eukaryotic genome [12] by permuting DNA segments according to their contact frequencies until the most likely scaffold was reached (see also [13]). Since these proof of concept studies, the assemblies of many genomes of various sizes from eukaryotes [e.g. 14–16] and prokaryotes [17] have been significantly improved using scaffolding approaches exploiting Hi-C data.

Although GRAAL was effective on medium-sized or small (<100 Mb) eukaryotic genomes such as that of the fungus *Trichoderma reesei* [18], scalability limitations were encountered when tackling genomes whose complexity and size required significant computer calculation capacity. Furthermore, as was also observed with other Hi-C-based scaffolders, the raw output of GRAAL includes a number of caveats that need to be corrected manually to obtain a finished genome assembly. To tackle these limitations, we developed instaGRAAL, an enhanced, open-source program optimized to reduce the computational load of chromosome scaffolding and that includes polishing steps to automatically complete the assembly process. The polishing, which aims to minimise assembly errors, can exploit available genetic linkage data.

InstaGRAAL was applied to the 214 Mb and 500 Mb haploid genomes of the common brown algae *Ectocarpus* sp. and *Desmarestia herbacea*, respectively. The *Ectocarpus* genome is currently only published in draft form [19] and the *Desmarestia* genome was previously unpublished. Brown algae are a group of complex multicellular eukaryotes that have been evolving independently from animal and land plants for more than a billion years. *Ectocarpus* sp. was the first species within the brown algal group to be sequenced, as a model organism to investigate multiple aspects of brown algal biology including the acquisition of multicellularity, sex determination, life cycle regulation and adaptation to the intertidal [20–23]. A range of genetic and genomic resources have been established for *Ectocarpus* sp. including a dense genetic map generated with 3,588 SNP markers [24], which was used to comprehensively validate the instaGRAAL assembly.

## Results

### From GRAAL to instaGRAAL

The core principles of GRAAL and instaGRAAL are similar: both exploit a MCMC approach to perform a series of permutations (insertions, deletions, inversions, swapping, etc.) of genome fragments (referred to here as a ‘bins’, see Methods) based on an expected contact distribution [12]. The parameters (A, *α* and δ) that describe this contact distribution are first initialized using a model inspired by polymer physics [25]. This model describes the expected contact frequency *P(s)* between two loci separated by a genomic distance *s* (when applicable):

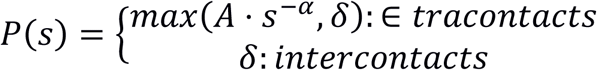

The parameters are then iteratively updated directly from the real scaffolds once their size increases sufficiently [12]. Each bin is tested in several positions relative to putative neighbouring fragments. The likelihood of each arrangement is assessed from the simulated or computed contact distribution, and the arrangement is either accepted or rejected [12]. This analysis is carried out in cycles, with a cycle being completed when all the bins of the genome have been processed in this way. Any number of cycles can be run iteratively and the process is usually continued until the genome structure ceases to evolve, as measured by the evolution of the parameters of the model. The core functions of the program use Python libraries as well as the CUDA programming language, and therefore necessitate a NVIDIA graphics card with at least 1 Gb of memory.

The technical limitations of GRAAL were i) high memory usage when handling Hi-C data for large genomes (*i.e.* over 100 Mb), 2) difficulties when installing the software, and 3) the need to adjust multiple *ad hoc* parameters to adapt to differences in genome size, read coverage, Hi-C contact distribution, specific contact features, etc. InstaGRAAL (https://github.com/koszullab/instaGRAAL) addresses all these shortcomings. First, we rewrote the memory-critical parts of the program, such as permutation sampling and likelihood calculation, so that they are computed using sparse contact maps. We reduced the software’s dependency footprint and added detailed documentation, deployment scripts and containers to ease its installation. Finally, we opened up multiple hard-coded parameters to give more control for end-users while improving the documentation on each of them, and selecting relevant default parameters that can be implemented for a wide range of applications (see options online, and Discussion). Overall, these upgrades result in a program that is lighter in resources, more flexible, and more user-friendly.

Other problems encountered with the original GRAAL program included 1) the presence of potential artefacts introduced by the permutation sampler, such as spurious permutations (e.g. local inversions) or incorrect junctions between bins; 2) difficulties with the correct integration of other types of data such as long reads and 3) difficulties with handling sequences that were either too short, highly repeated or with low coverage. We addressed these points by identifying and putting aside these problematic sequences during a filtering step. These sequences are subsequently reinserted into the final scaffolds, whenever possible (see Methods), with the help of linkage data when available. Overall, when compared to the raw GRAAL output, the resulting “polished” assemblies were significantly more complete and more faithful to the actual chromosome structure.

### Scaffolding of the *Ectocarpus* sp. chromosomes with instaGRAAL

To test and validate instaGRAAL, we generated an improved assembly of the genome of the model brown alga *Ectocarpus* sp.. A reference genome consisting of 1,561 scaffolds generated from Sanger sequence data is available [20]. A Hi-C library was generated from a clonal culture of a haploid partheno-sporophyte carrying the male sex chromosome using a GC-neutral restriction enzyme (DpnII). The library was paired-end sequenced (2×75 bp – the first ten bases were used as a tag and to remove PCR duplicates) on a NextSeq apparatus (Illumina). Of the resulting 80,521,968 paired-end reads, 41,288,678 read pairs were aligned unambiguously along the reference genome using bowtie2 (quality scores below 30 were discarded), resulting in 2,554,639 links bridging 1,806,386 restriction fragments (Fig 1a) (see Methods for details on the experimental and computational steps). The resulting contact map in sparse matrix format was then used to initialize instaGRAAL along with the restriction fragments (RFs) of the reference genome (Fig 1a-b) (see Table S1 for an example of sparse file matrix).

**Fig. 1 :**
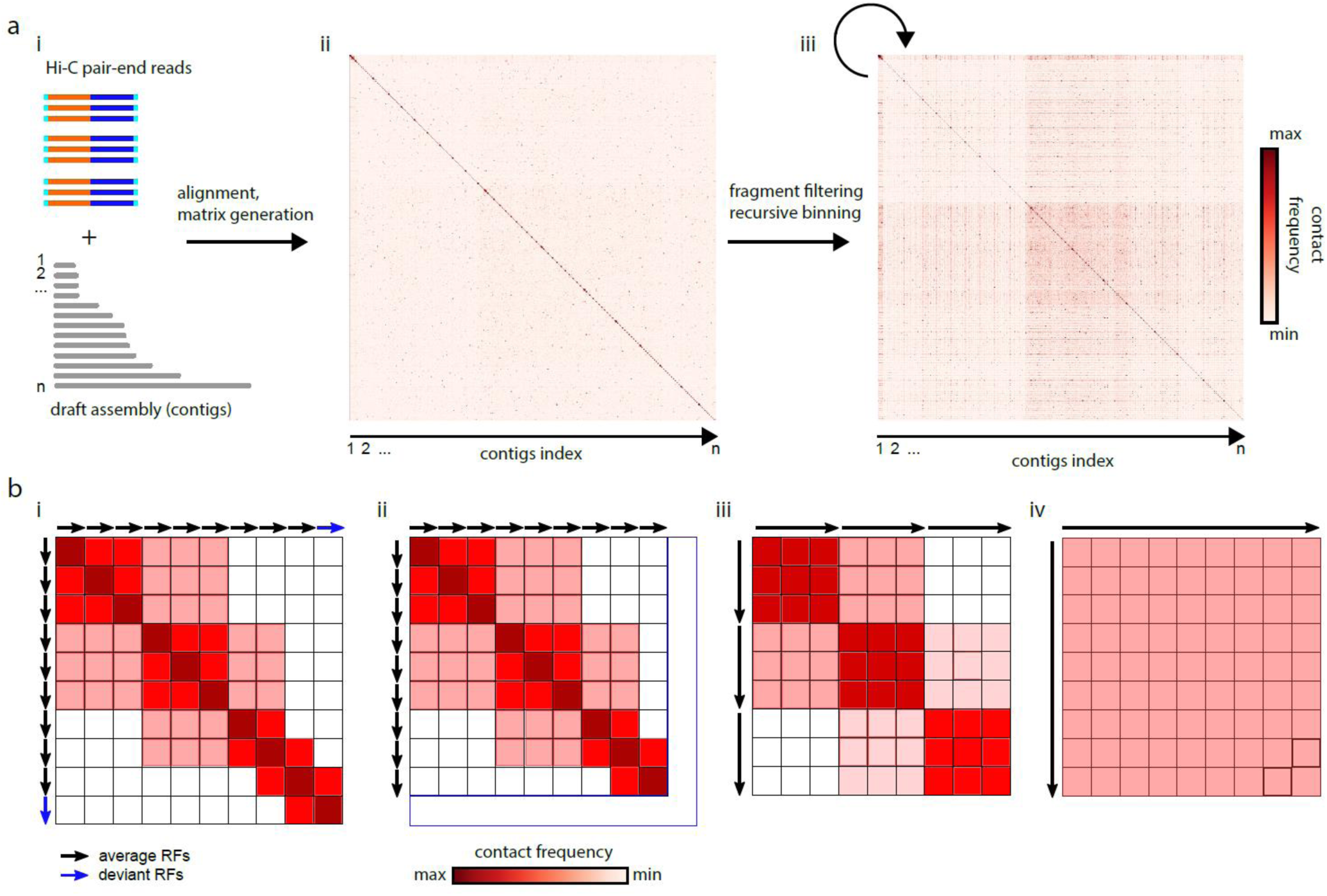
Matrix generation and binning process. (a) (from left to right): i) The input data to be processed, paired-end reads to be mapped onto the *Ectocarpus*. sp. draft assembly; ii) Raw contact map before binning: each pixel is a contact count between two restriction fragments (RF); iii) Raw contact map after binning: each pixel is a contact between a determined numbers of RFs (see b). (b) Schematic description of one iteration of the binning process over 10 restriction fragments (arrows). From left to right: i) initial contact map, each pixel is a contact count between two RFs; ii) filtering step: RFs either too short or presenting a read coverage below one standard deviation below the mean are discarded; iii) binning step (1 bin = 3RFs): adjacent RFs are pooled by three, with sum-pooling along all pixels in a 3×3 square.

### Convergence of the *Ectocarpus* sp. assembly towards 27 major scaffolds

Given the probabilistic nature of the algorithm, we evaluated the program’s consistency by running it three times with different resolutions. Briefly, we filtered out RFs that were shorter than 50 bp and/or whose coverage was one standard deviation below the mean coverage. Then, we sum-pooled (or binned) the sparse matrix by groups (or bins) of three RFs five times, recursively (Fig 1a-b). Each recursive instance of the sum-pooling is subsequently referred to as a level of the contact map. A level determines the resolution at which permutations are being tested: the higher the level, the lower the resolution, the longer the sequences being permuted and, consequently, the faster the computation. The binning process is shown in Fig. 1b. Regarding *Ectocarpus* sp., we found that level four (bins of 81 RFs) was an acceptable balance between high resolution and fast computation on a desktop computer with a GeForce GTX TITAN Z graphics card. Moreover, whether instaGRAAL was run at level four, five or six (equivalent to bins of 81, 243 and 729 RFs respectively), all assemblies quickly (∼6hrs) converged towards similar genome structures (Fig. 2a).

**Fig. 2 :**
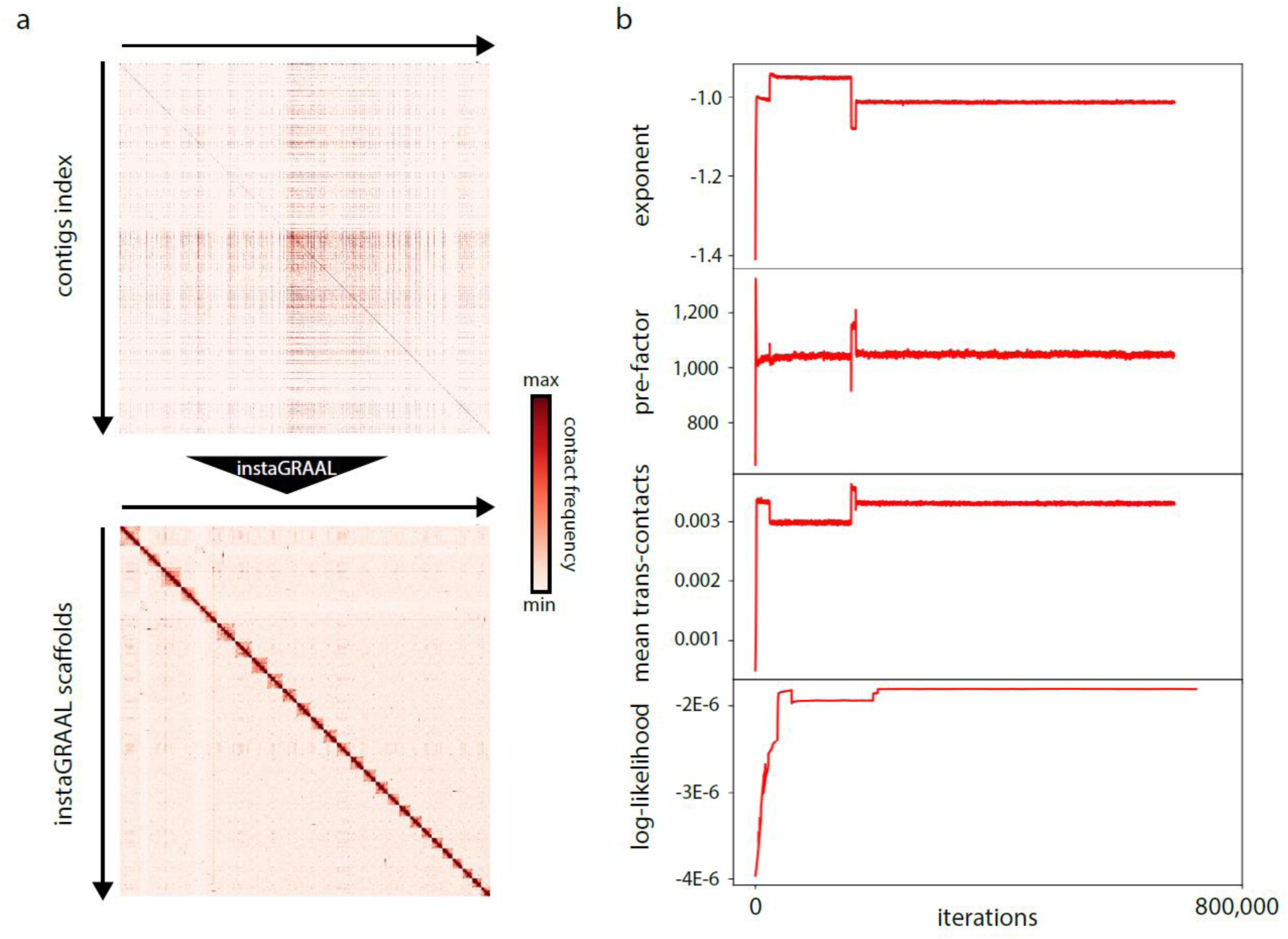
Evolution of the *Ectocarpus* sp. contact map, the parameters of the polymer model and the log-likelihood of the contact map. (a) The raw contact map before (upper part) and after (bottom part) scaffolding using instaGRAAL. Scaffolds are ordered by size. (b) Evolution of three parameters of the polymer model (exponent, pre-factor, mean trans-contacts) and the log-likelihood through the iterations.

We plotted the evolution of the log-likelihood and of model parameters as a function of the number of iterations (Fig. 2b). The interquartile ranges (IQR, used to indicate stability in Marie-Nelly et al., 2014 [12]) of all parameters decreased to near-zero values at the end of each scaffolding run, indicating that they all stably converged and that the final structures oscillated near the final values in negligible ways. More qualitatively, each run led to the formation of 27 main scaffolds (Fig. 2a) with the 27^th^ largest scaffold being more than a hundred times longer than the 28^th^ largest one (Fig. 3) (movie S1). Each of the 27 scaffolds was between four and ten times longer than the combined length of the remaining sequences (Fig. 3). This strongly suggests that the 27 scaffolds correspond to chromosomes, a number consistent with karyotype analyses [26]. Taken together, these results indicate that instaGRAAL successfully assembled the *Ectocarpus* sp. genome into chromosome-level scaffolds. As the supplementary movie suggests, scaffold-level convergence is visible after only a few cycles, indicating that instaGRAAL is able to quickly determine the global genome structure most likely to fit the contact data. The remainder of the cycles are devoted to intra-chromosomal refinement.

**Fig. 3:**
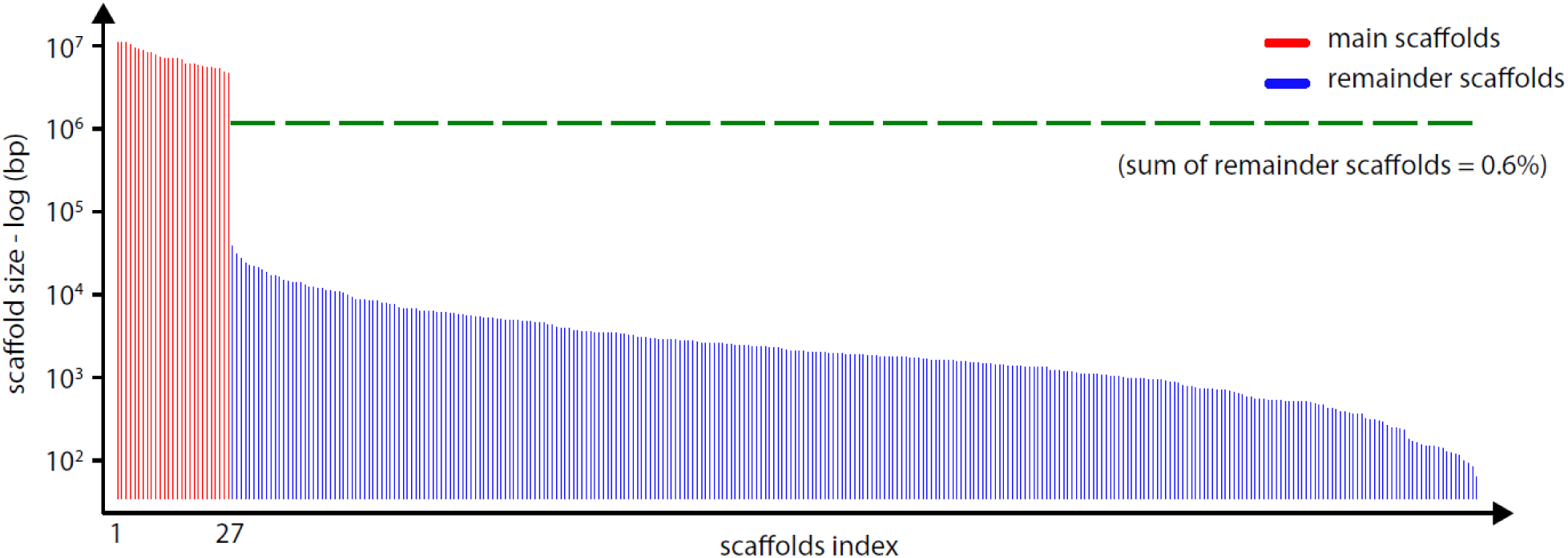
Size distribution (log scale) of the final *Ectocarpus* sp. scaffolds after 250 instaGRAAL iterations. After filtering, and prior polishing, 27 main scaffolds (red bars) or putative chromosomes were obtained. The dotted green horizontal line represents the proportion of the filtered genome that was not integrated into the main 27 scaffolds and represent less than 0.6% of the initial assembly. Each scaffold presents, after normalization, a high quality Hi-C profile with features that are typical of eukaryotic genomes (Figure S2).

### Polishing the chromosome-level assembly of the *Ectocarpus* sp. genome

InstaGRAAL also includes a number of procedures that aim to correct modifications of the reference assembly contigs introduced during the Hi-C scaffolding (Fig. 4).

**Fig. 4:**
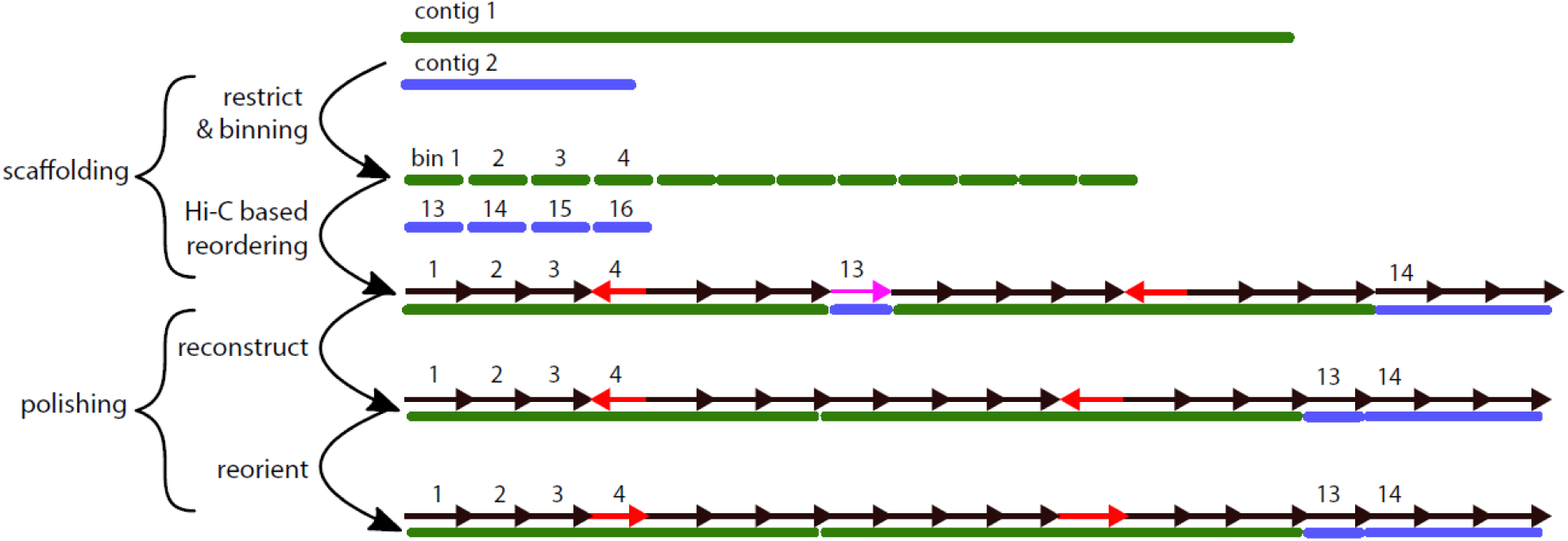
Step-by-step correction procedure. Polishing procedure (top to bottom): i) *in silico* restriction of the genome and binning, yielding a set of bins; ii) reordering of all bins into scaffolds without reference to their contig of origin; typically, groups of bins from the same contig naturally aggregate, but some bins get scattered to other scaffolds (e.g. bin 13, pink arrow), while others will be “flipped” with respect to the original assembly (e.g. bin 4, red arrows); iii) reconstruction of the original contigs by relocating scattered bins next to the biggest bin group; iv) bins in the original contigs are oriented according to their original consensus orientation.

These modifications principally involve discrete inversions or insertions of DNA segments (typically corresponding to single bins or RFs) (see also [12]). Such alterations are inherent to the statistical nature of instaGRAAL, which will occasionally improperly permute neighbouring bins because of the high density of contacts between them. These are part of a broader set of assembly errors (subsequently referred to as ‘misassemblies’) that we detected by aligning the reference contigs with the instaGRAAL scaffolds and analysing the mapping results using QUAST. We corrected these errors as follows: first, all bins processed by instaGRAAL that belonged to the same contig were constrained to their original orientation (Fig 4). If a contig was split across multiple scaffolds, the smaller parts of this contig were relocated to the largest one, respecting the original order and orientation of the bins. Then we reinserted whenever possible sequences that had been filtered out prior to instaGRAAL processing (e.g. contig extremities with poor read coverage; see Methods and Marie-Nelly et al., 2014 [12]) into the chromosome level scaffold at their original position in the contig of origin. A total of 3,832,980 bp were reinserted into the assembly this way. These simple steps alleviated artificial contig truncations observed with the original GRAAL program.

Some filtered bins had no reliable region to be associated with post scaffolding, because the entire initial contig they belonged to had been filtered. These sequences, which were left as-is and appended at the end of the genome, consisted of 543 scaffolds spanning 3,141,370 bp, i.e. < 2% of the total DNA. Together, these steps removed all the misassemblies detected by QUAST (Table 1).

**Table 1.** Comparison of Nx, NGx (Nx with respect to the reference; in bp) stats and other genomics statistics for the different assemblies (Pseudochromosomes, GRAAL and instaGRAAL) of the *Ectocarpus* sp. genome.

| Assembly | Reference genome | Pseudochromosomes | GRAAL | instaGRAAL |
| --- | --- | --- | --- | --- |
| N50 | 497,380 | 6,528,661 | 6,867,074 | 6,813,345 |
| NG50 | 497,380 | 6,528,661 | 6,725,743 | 6,813,345 |
| N75 | 233,412 | 5,613,161 | 5,693,784 | 5,686,617 |
| NG75 | 233,412 | 5,613,161 | 5,672,622 | 5,686,617 |
| L50 | 118 | 12 | 11 | 11 |
| LG50 | 118 | 12 | 12 | 11 |
| L75 | 258 | 19 | 18 | 19 |
| LG75 | 258 | 19 | 19 | 19 |
| # genomic features | 362,921 | 350,497 + 7,261 part | 342,253 + 9,766 part | 350,555 + 7,261 part |
| Complete BUSCO (%) | 75.9 | 76.9 | 76.24 | 77.56 |
| K-mer-based compl. (%) | 100 | 99.97 | 98.53 | 100 |

To further validate the assembly, we exploited genetic linkage data to search for potential translocations between scaffold extremities [24]. This analysis, now implemented as an option in instaGRAAL, detected such events in the unpolished version but none in the polished assembly. The polished instaGRAAL assembly is therefore fully consistent with the genetic recombination data, confirming the efficiency of the approach.

### Comparisons with previous *Ectocarpus* sp. genome assemblies and validation of the instaGRAAL assembly

We compared the polished instaGRAAL assembly (subsequently referred to as the polished assembly) with three earlier assemblies of the *Ectocarpus* sp. genome (Table1 and Table S2): 1) the reference assembly generated using Sanger sequencing data [20], mentioned above, which was assumed to be highly-accurate but highly fragmented (1,561 scaffolds); 2) an assembly generated by combining genetic recombination data and the Sanger assembly [19,24] (pseudochromosome assembly) and 3) an assembly generated by running the original GRAAL program on the reference genome (GRAAL assembly).

We aligned the instaGRAAL, pseudochromosomal and GRAAL assemblies to the reference assembly to detect misassemblies and determine whether the genome annotations (362,919 features) were conserved. We then validated each assembly using genetic linkage data (Methods). For each assembly, we computed the following metrics: the number of misassemblies, the fraction of conserved annotations, ortholog completeness and cumulative length/Nx distributions (Table 1). These assessments were carried out using BUSCO [27] for orthologue completeness (Figure S1) and QUAST-LG’s validation pipeline [28] for the other tests. QUAST-LG is an updated version of the traditional QUAST pipeline specifically designed for large genomes. We followed the terminology used by both programs, such as the BUSCO definition of ortholog and completeness, as well as QUAST’s classification system of contig and scaffolds misassemblies.

The polished instaGRAAL assembly was of better quality than both the pseudochromosome and GRAAL assemblies (Table 1 and Figure S1). The polished assembly incorporated 795 of the reference genome scaffolds (96.8% of the sequence data) into the 27 chromosomes based on the high density genetic map [19], compared to 531 for the pseudochromosomal assembly (90.5% of the sequence data). Moreover, this assembly contained fewer misassemblies, retained more annotations and was more complete (both in terms of k-mers and BUSCO ortholog content). For some metrics the differences were marginal, but always in favour of the instaGRAAL assembly. BUSCO completeness was similar (76.2%, 76.9%, 77.6% for the GRAAL, pseudochromosomal and instaGRAAL assemblies, respectively, Figure S1), and an improvement over the 75.9% of the reference. These absolute numbers remain quite low, presumably because of the lack of a set of orthologs well adapted to brown algae.

All quantitative metrics, such as N50, L50 and cumulative length distribution, increased dramatically when compared with the reference genome (Table 1). N50 increased more than tenfold, from 496,777 bp to 6,867,074 bp after the initial scaffolding, and to 6,942,903 bp after the polishing steps. Ninety nine point four percent of DNA sequence of the 1,018 contigs was integrated into the 27 largest scaffolds after instaGRAAL processing. Overall, the analysis indicated that many of the rearrangements found in the pseudochromosome assembly were potentially errors, and that both GRAAL and instaGRAAL were efficient at placing large regions where they belong in the genome, albeit less accurately for GRAAL and in the absence of polishing. These statistics underline the importance of the post-scaffolding polishing steps, and the usefulness of a program that automates these steps.

### Comparison between the *Ectocarpus* sp. instaGRAAL and pseudochromosomal assemblies

Compared to the pseudochromosomal assembly, the instaGRAAL assembly lost 23 scaffolds but gained 287 that the genetic map was unable to anchor to chromosomes (Table S2). We observed few conflicts between the two assemblies. One major difference is that instaGRAAL was able to link the 4^th^ and 28^th^ pseudochromosomes that were considered to be separate by the genetic map [24] because of the limited number of recombination events observed. The fusion in the instaGRAAL assembly is consistent with the fact that the 28^th^ pseudochromosome is the smallest of the linkage groups, with only 54 markers over 41.8 cM and covering 3.8 Mb. The 28^th^ pseudochromosome has a very large gap which might reflect uncertainty in the ordering of the markers. Interestingly, this gap is located at one end of the linkage group, precisely where instaGRAAL now detects a fusion with the 4^th^ pseudochromosome. In addition, the fact that there is no mix between the 4^th^ and 28^th^ pseudochromosomes on the merged instaGRAAL chromosome but rather a simple concatenation, suggests that the genetic map was unsuccessful in joining those two linkage groups, but that instaGRAAL correctly assembled the two pseudochromosomes (see Table S3 for correspondences between pseudochromosomes and instaGRAAL super scaffolds). InstaGRAAL was also more efficient than the genetic map in orienting scaffolds (Table S2). Among the scaffolds that were oriented in the pseudochromosomal assembly, about half of the ‘plus’ orientated were actually ‘minus’ and vice versa. The limited number of markers detected in the scaffolds anchored to the genetic map was likely the reason for this high level of incorrect orientations.

### Scaffolding of the *Desmarestia herbacea* genome

To test and validate instaGRAAL on a second, larger genome, we generated an assembly of the haploid genome of *D. herbacea*, a brown alga that had not been sequenced before. We set up the assembly pipeline and subsequent scaffolding from raw sequencing reads to assess the robustness of instaGRAAL with *de novo*, non-curated data. The pipeline proceeded as follows: first, we acquired 259,556,174 short paired-end shotgun reads (Illumina HiSeq2500 and 4000) as well as 1,353,202 long reads generated using PacBio and Nanopore (about 150X short reads and 15X long reads). Sequencing reads were processed using the hybrid MaSuRCA assembler (v3.2.9) [29], yielding 7,743 contigs representing 496 Mb. We generated Hi-C data following a protocol similar to that used for *Ectocarpus* sp. (Material and Methods). Briefly, 101,879,083 reads were mapped onto the hybrid assembly, yielding 7,649,550 contacts linking 1,359,057 fragments. We then ran instaGRAAL using similar default parameters to that used for *Ectocarpus* sp., for the same number of cycles. Polishing was applied to the resulting scaffolds. The scaffolding process resulted in 40 scaffolds larger than 1 Mb (Figure S3), representing 98.1% of the initial, filtered scaffolding and 89.3% of the total initial genome after polishing and reintegration. The exact number of chromosomes in *D. herbacea* is unknown, with estimations varying between ∼23, and possibly up to 29 [30]. Most (35; Table S4) of the scaffolds generated by instaGRAAL were syntenic with the 27 *Ectocarpus* sp. scaffolds with an average of 61% orthologous conservation. Among the remaining five scaffolds, one (#7571, 4.6Mb) corresponded to a bacterial genome of the *Planctomycetes* clade, a species typically present in algal culture. Two other large scaffolds (#7565, 4.4Mb and #5576, 2.8Mb) annotated as *Bacteroidetes* displayed highly divergent GC content (37 and 40% vs. 48% for the rest of the genome) and no *Desmarestia* gene prediction, suggesting that we captured the DNA from bacteria species present in the *Desmarestia* culture tanks. One scaffold (#7563, 1.1Mb) was annotated as viral DNA and could reflect the insertion of a Pheaovirus in the *Desmerastia* genome, similarly to insertion of EsV1 genome in *Ectocarpus sp*.

Overall, instaGRAAL successfully scaffolded most of the *Desmarestia* genome. Although the final number of scaffolds remained slightly higher that the estimated number of chromosomes in this species, visual inspection of the scaffolds did not suggest obvious large scale scaffolding mistakes. Comparative analysis with *Ectocarpus sp*. and other species currently being sequenced will further refine this analysis, and improve the assembly.

### Comparisons with existing methods

To date, only a limited number of Hi-C based scaffolding programs are publicly available, and, as far as we can tell, no detailed comparison has been performed between the existing programs to assess their respective qualities and drawbacks. In an attempt to benchmark instaGRAAL, we ran SALSA2 [31] and 3D-DNA on the same *Ectocarpus* sp. and *Desmarestia herbacea* reference genome and Hi-C reads. 3D-DNA is a scaffolder that was hallmarked with the assembly of *Aedes aegypti* and SALSA2 is a recent program with a promising approach that directly integrates Hi-C weights into the assembly graph. For *Ectocarpus* sp., SALSA2 ran for nine iterations and yielded 1,042 scaffolds, with an N50 of 6,552,506 (L50 = 11). Its BUSCO-completeness was 77.6%, a level identical to that obtained with instaGRAAL. Overall the metrics were satisfactory but SALSA2 was outperformed by instaGRAAL post-polishing. The contact map of the resulting SALSA2 assembly, displayed noticeably unfinished scaffolds (Figure S4 and S5). This, coupled with a lower N50 value, suggests that instaGRAAL is more successful at merging scaffolds when appropriate.

We computed similar size and completeness statistics for the final instaGRAAL *D. herbacea* assembly and compared these to the values obtained with SALSA2 and 3D-DNA. We also mapped the Hi-C reads onto all three final assemblies in order to qualitatively assess the chromosome structure. The results are summarized in Table S5.

Briefly, statistics across assemblies were similar, the polished instaGRAAL assembly had 73% BUSCO completeness, consistent with the values of 73.6% and 70.3% obtained for SALSA2 and 3D-DNA, respectively. However, the Lx/Nx metrics diverged significantly, the instaGRAAL assembly N50 was 12.4 Mb, similar to SALSA2 (12.8) and much larger than 3D-DNA (0.2 Mb). However, visual inspection of the contact maps indicated that neither SALSA2 nor 3D-DNA succeeded in fully scaffolding the genome of *Desmarestia herbacea* (Figure S4). Notably, SALSA2 created a number of poorly-supported junctions to generate chromosomes, whereas 3D-DNA failed to converge towards any kind of structure. In contrast, although the instaGRAAL final assembly still contains contigs that are incorrectly positioned, a coherent structure corresponding to 40 scaffolds (including contaminants) emerged (Figure S4). One possibility is that the *de novo* MaSuRCA assembly was low-quality, which could have resulted in alignment errors that disrupted the contact distribution and subsequent Hi-C scaffolding. Another possible explanation for these differences is that it remains difficult to dissect all the options and tuneable parameters of these scaffolders, and therefore that we did not find the optimal combination with respect to the *D. herbacea* draft assembly. Nevertheless, these results highlight the robustness of instaGRAAL which was able to successfully scaffold the *D. herbacea* genome using default parameters.

## Discussion

InstaGRAAL is a Hi-C scaffolding program that can process large eukaryotic genomes. Below we discuss the improvements made to the program, its remaining limitations and the steps that will be needed to tackle them.

### Reference-based polishing

An important improvement of instaGRAAL compared to GRAAL relates to post-scaffolding polishing. Local misassemblies, e.g. local bin inversions or disruptive insertions of small scaffolds within larger ones, are an inevitable consequence of the algorithm’s most erratic random walks. These small misassemblies are retained because flipping a bin doesn’t markedly change the relative distance of an RFs relative to its neighbours, and because small scaffolds typically carry less signal and therefore exhibit a greater variance in terms of acceptable positions. Depending on the trust put in the initial set of contigs, one may be unwilling to tolerate these changes as well as “partial translocations”, i.e. the splitting of an original contig into two scaffolds. The prevalence of such mistakes can be estimated by comparing the orientation of bins relative to their neighbours in the instaGRAAL assembly vs. the original assembly. Our assumption is that, if a single bin was flipped or split by instaGRAAL, this was likely a mistake that needed to be corrected by the polishing procedures. Consequently, we chose to remain faithful to the reference contigs, given that the initial *Ectocarpus* sp. reference was based on very accurate Sanger reads. Polishing therefore aims at reinstalling the initial contig structure and orientation while preserving to a maximum extent the overall instaGRAAL scaffold structure.

In addition, polishing reintegrates into the assembly the bins removed during the initial filtering process according to their position along the original assembly contigs. Most filtered bins corresponded to the extremities of the original contigs, because their size depended on the position of the restriction sites within the contig, or because they consisted of repeated sequences with little or no read coverage. The tail filtering polishing step inserts these bins back at the extremities of these contigs in the instaGRAAL assembly.

The combination of a probabilistic algorithm with a deterministic polishing step provides robustness to instaGRAAL. First, the MCMC step identifies, with few prior assumptions, a high-likelihood family of genome structures, almost always very close to the correct global scaffolding. The polishing step combines this result with prior assumptions made about the initial contig structures generated through robust, established assembly programs, refining the genomic structure within each scaffold. To give the user a fine-grained degree of control over polishing, the implementation itself into instaGRAAL is split into independent modules that each make an assumption about the initial contig structure necessary to perform the correction: the ‘reorient’ module assumes that the initial contigs do not display inversions, the ‘rearrange’ module assumes that there are no relocations within contigs.

We would like to underline that, despite the improvements brought about by these new procedures, instaGRAAL assemblies remain perfectible, notably because of the reliance on the quality of the original contig assembly. However, chromosome-level assemblies consistent with known genetic maps and other datasets were reached every time the program was applied on a new species.

### Sparse data handling

The implementation of a sparse data storage method in instaGRAAL allows much more intense computation than with GRAAL. Because the majority of map regions are devoid of contacts, instaGRAAL essentially halves the order of magnitude of both algorithm complexity and memory load, i.e. they increase linearly with the size of the genome instead of geometrically. This improvement potentially allows the assembly of Gb-sized genomes in five to six days using a desktop computer (and faster with a larger computational resource).

### Filtering

Variations in GC% along the genome, and/or other genomic features, can lead to variation in Hi-C read coverage and impair interpretation of the Hi-C data. Correction and attenuation procedures that alleviate these biases are therefore commonly used in Hi-C studies [32– 34]. However, these procedures are not compatible with instaGRAAL’s estimation of the contact distribution (for more see [35]). A subset of bins will therefore diverge strongly from the others, displaying little if no coverage. A filtering step is needed to remove these bins as they would otherwise impact the contact distribution and the model parameter estimation. These disruptive bins represent a negligible fraction of the total genome (< 3% of the total genome size of *Ectocarpus* sp., for instance) and are reincorporated into the assembly during polishing. On the other hand a subset of bins representing small, individual scaffolds are not reinserted during polishing, and are added to the final assembly as extra-scaffolds (as in all sequencing projects). Additional analyses and new techniques such as long or linked reads are needed to improve the integration of these scaffolds into the genome.

### Resolution

The binning procedure will influence the structure of the final assembly as well as its quality. For example, low level binning (e.g. one bin = three RFs) will lead to an increased number of bins and a large, sparse contact map with a low signal-to-noise ratio, where many of the bins display poor read coverage as on average they will have fewer contacts with their immediate neighbours. Because of the resulting low signal-to-noise ratio, an invalid prior model will be generated and, when referring to this model, the algorithm will fail to scaffold the bins properly, if at all. Moreover, due to its probabilistic nature, the algorithm will generate a number of false positive structural modifications such as erroneous local inversions or permutations of bins. The numerous bins will create more genome structures to explore to handle all the potential combinations, and exploring this space until convergence will take longer and be computationally demanding.

An optimal resolution is therefore a compromise between the bin size, the coverage, and the quality of the original contig assembly. Although a machine powerful enough operating on an extremely contact-rich matrix would be successful at any level, it is unclear whether such resources are necessary. Our present assemblies (e.g. 1 bin = 81 RFs for both; Material and Methods) had good quality metrics after a day’s worth of calculation on a standard desktop computer. Moreover convergence was qualitatively obvious after a few cycles. This suggests that more computational power yields diminishing returns, and therefore that appropriate polishing is a more efficient approach to correct any remaining misassemblies.

### Binning

The fragmentation of the original assembly used to generate the initial contact map has obviously a substantial effect on the quality of the final scaffolding. Because binning cannot be performed beyond the resolution of individual contigs, however small they may be, there is a fixed upper limit to the a scale at which a given matrix can be binned. A highly fragmented genome with many small contigs will necessarily generate a high-noise, high-resolution matrix. Attempts to reassemble a genome based on such a matrix will run into the problems discussed above (resolution). This limitation can be alleviated, to some extent, by discarding the smallest contigs, with the hope that the remaining contigs will cover enough of the genome. The contigs that are removed can be reintegrated into the assembly during the polishing steps. This ensures an improved Nx metric while retaining genome completeness. It should be noted, however, that the size of the contigs is only important insofar as they need to contain sufficient restriction sites, and each of the restriction fragments must have sufficient coverage. The choice of enzyme and the frequency of its corresponding site is thus crucial. For instance, with an average of one restriction site every 600 to 1,000 bp for *Dpn*II, contigs as short as 10 kb may contain enough information to be correctly reassembled. The restriction map therefore strongly influences both the minimum limit on N50 and genome fragmentation.

### Benchmarking

In order to test our tool against existing programs, we ran two scaffolders available online (SALSA2 and 3D-DNA) on our two genomic datasets. In all instances, instaGRAAL proved more successful at scaffolding both genomes. However, we have not extensively tested all the combinations of parameters of both programs, and acknowledge the difficulty in designing and implementing Hi-C scaffolding pipelines with extensive dependencies that compound the initial complexity of the task and add yet more configurable options to know in advance. Finding the correct combination of CUDA and Python dependencies to install instaGRAAL on a given machine can be challenging as well. Therefore, our benchmarking attempt should be rather seen as a way to stress the importance of implementing sensible default parameters that readily cover as many use cases as possible for the end user. There is almost no doubt that both 3D-DNA and SALSA2, with the appropriate parameters and polishing steps, would produce satisfying scaffolding; on the other hand, knowing which input parameters has to be specified in advance is a nontrivial task, especially given the computational resources needed for a single scaffolding run. With instaGRAAL, we wish to combine the simplicity of a default configuration that works in most instances, with the flexibility offered by the power of MCMC methods.

### Integrating information from the Hi-C analysis with other types of data

Aggregating data from multiple sources to construct a high-quality assembly remains a challenging problem with no systematic solution. As long read technologies become more affordable, there is an increasing demand to reconcile the scaffolding capabilities of Hi-C based methods with the ability of long reads to span regions that are difficult to assemble, such as repeated sequences. The most intuitive approach would be to perform Hi-C scaffolding on an assembly derived from high-coverage and corrected long reads, as was done for several previous assembly projects [14,36]. Alternative approaches also exist, such as generating Hi-C and long-read-based assemblies separately and merging them using programs such as CAMSA (Aganezov et al., 2017) or Metassembler (Wences et al., 2015). Lastly, pipelines such as PBJelly (English et al., 2012) have proven successful at filling existing gaps in draft genomes, regardless of their origin, with the help of long reads. InstaGRAAL shows that high quality metrics can still be attained without the help of long reads, but it can nevertheless integrate them when necessary or available. Long reads are not the only type of data that may be used to improve assemblies. Linkage maps, RNA-seq, optical mapping and 10X technology all provide independent data sources that can help improve genome structure and polish specific regions. The success of future assembly projects will hinge on the ability to process these various types of data in a seamless and efficient manner.

## Material and Methods

### Preparation of the Hi-C libraries

The Hi-C library construction protocol was adapted from [7,37]. Briefly, partheno-sporophyte material was chemically cross-linked for one hour at RT using formaldehyde (final concentration: 3% in 1X PBS; final volume: 30 ml; Sigma-Aldrich, St. Louis, MS). The formaldehyde was then quenched for 20 min at RT by adding 10 ml of 2.5 M glycine. The cells were recovered by centrifugation and stored at −80°C until use. The Hi-C library was then prepared as follow. Cells were resuspended in 1.2 mL of 1X *Dpn*II buffer (NEB, Ipswich, MA), transferred to a VK05 tubes (Precellys, Bertin Technologies, Rockville, MD) and disrupted using the Precellys apparatus and the following program ([20 sec – 6000 rpm, 30 sec – pause] 9x cycles). The lysate was recovered (around 1.2 mL) and transferred to two mL tubes. SDS was added to a final concentration of 0.3% and the 2 reactions were incubated at 65°C for 20 minutes followed by an incubation of 30 minutes at 37°C. A volume of 50 µL of 20% triton-X100 was added to each tube and incubation was continued for 30 minutes. *Dpn*II restriction enzyme (150 units) was added to each tube and the reactions were incubated overnight at 37°C. Next morning, reactions were centrifuged at 16,000 x g for 20 minutes. The supernantants were discarded and the pellets were resuspended in 200 µL of NE2 1X buffer and pooled (final volume = 400 µL). DNA extremities were labelled with biotin using the following mix (50 µL NE2 10X buffer, 37.5 µL 0.4 mM dCTP-14-biotin, 4.5 µL 10mM dATP-dGTP-dTTP mix, 10 µL Klenow 5 U/µL) and an incubation of 45 minutes at 37°C. The labelling reaction was then split in two for the ligation reaction (ligation buffer – mL, ATP 100 mM – 160 µL, BSA 10 mg/mL – 160 µL, ligase 5 U/µL – 50 µL, H2O – 13.8 mL). The ligation reactions were incubated for 4 hours at 16°C. After addition of 200 µL of 10%, SDS 200 µL of 500 mM EDTA and 200 µL of proteinase K 20 mg/mL, the tubes were incubated overnight at 65°C. DNA was then extracted, purified and processed for sequencing as previously described (Lazar-Stefanita et al., 2017). Hi-C libraries were sequenced on a NextSeq 550 apparatus (2 × 75 bp, paired-end Illumina NextSeq with the first ten bases acting as barcodes; Marbouty et al., 2014).

### Contact map generation

Contact maps were generated from reads using the hicstuff pipeline for processing generic 3C data, available at https://github.com/koszullab/hicstuff. The backend uses the bowtie2 (version 2.2.5) aligner run in paired-end mode (with the following options: --maxins 5 –very-sensitive-local). Alignments with mapping quality lower than 30 were discarded. The output was in the form of a sparse matrix where each fragment of every chromosome was given an unique identifier and every pair of fragments was given a contact count if it was nonzero. Fragments were then filtered based on their size and total coverage. First, fragments shorter than fifty base pairs were discarded. Then, fragments whose coverage was less than one standard deviation below the mean of the global coverage distribution were removed from the initial contact map. A total of 6,974,350 bp of sequence was removed this way. An initial contact distribution based on a simplified a polymer model [25] with three parameters was first computed for this matrix. Finally, the instaGRAAL algorithm was run using the resulting matrix and distribution.

For the *Ectocarpus* sp. genome, instaGRAAL was run at level 4 (n = 81 RFs), 5 (n = 243 RFs) and 6 (n = 729 RFs). Levels 5 and 6 were only used to check for genome stability and consistency in the final chromosome count. Level 4 was used for all subsequent analyses. All runs were performed for 250 cycles. The starting fragments for the analysis were the reference genome entirely fragmented into restriction fragments. The MCMC was run with 3 burn-in cycles. The same parameters were used for the *Desmarestia herbacea* genome.

### Polishing of genome assemblies

The assembled genome generated by instaGRAAL was polished to remove misassemblies using a number of simple procedures that aimed to reinstate the local structure of the initial contigs where possible. Briefly, bins belonging to the same initial contig were juxtaposed in the same relative positions as in the starting assembly contig. Small groups of bins were preferentially moved to the location of larger groups when several such groups were present in the assembly. The orientations of sets of bins that had been regrouped in this manner were modified so that orientation was consistent and matched that of the majority of the group, re-orientating minority bins when necessary. Both steps are illustrated in Fig. 4. Finally, fragments that had been removed during the filtering steps were reincorporated if they had been adjacent to an already integrated bin in the initial assembly. The remaining sequences that could not be reintegrated this way were appended as non-integrated scaffolds.

### Validation metrics

Initial and final assembly metrics (Nx, GC distribution) were obtained using QUAST-LG [28]. Misassemblies were quantified using QUAST-LG with the minimap2 aligner in the back-end. Ortholog completeness was computed with BUSCO (v3) [27]. Assembly completeness was also assessed with BUSCO. The evolution of genome metrics between cycles was obtained using instaGRAAL’s own implementation.

### Validation with the genetic map

The validation procedure with respect to linkage data was implemented as part of instaGRAAL. Briefly, the script considers a set of pseudochromosomes where regions are separated by SNP markers, and a set of Hi-C scaffolds where regions are bins separated by restriction sites. It then finds best-matching pairs of pseudochromosomes/scaffolds by counting how many of these regions overlap from one set to the other. Then, for each pair, the bins in the Hi-C scaffold are rearranged so that their order is consistent with that of the corresponding pseudochromosome. Such rearrangements are parsimonious and try to alter as little as possible. Since there isn’t a one-to-one mapping from restriction sites to SNP markers, some regions in the Hi-C scaffolds are not present in the pseudochromosomes, in which case they are left unchanged. When the Hi-C scaffolds are altered this way, as was found in the case of the raw GRAAL assembly, the script acts as a correction. When the scaffolds are unchanged, as was the case with the instaGRAAL assembly, the script acts as a validation.

### Benchmarking

For each genome, the 3D-DNA program was run using the run-assembly-pipeline.sh entrypoint script with the following options: -i 1000 --polisher-input-size 10000 --splitter-input-size 10000. The Hi-C data was prepared with the Juicer pipeline as recommended by 3D-DNA’s documentation. The SALSA2 program was run with the –cutoff=0 option, and misassembly correction with the –clean=yes option. No expected genome size was provided. The program halted after 9 iterations for *Ectocarpus* sp., and 18 iterations for *Desmarestia herbacea*. Hi-C data was prepared with the Arima pipeline as recommended by SALSA2’s documentation.

### Software tool requirements

The instaGRAAL software is written in Python 3 and uses CUDA for the computationally intensive parts. It requires a working installation of CUDA with the pycuda library. CUDA is a proprietary parallel computing framework developed by NVIDIA, and requires a NVIDIA graphic card. The scaffolder also requires a number of common scientific Python libraries specified in its documentation. The instaGRAAL website lists computer systems onto which the program was successfully installed and run.

## List of abbreviations

RF: Restriction fragment
MCMC: Markov Chain Monte Carlo
LD: Linkage desiquilibrium
IQR: Inter-quartile range
3C: chromosome conformation capture
GRAAL -3D: genome (re)assembly assessing likelihood from 3D

## Availability of data and materials

The raw, unfiltered datasets generated and analysed during the current study are available in the SRA repository, SRR8550777.

The instaGRAAL software and its documentation are available at https://github.com/koszullab/instaGRAAL.

Assemblies, contact maps and relevant materials for the reproduction of the main results and figures are available at https://github.com/koszullab/ectocarpus_scripts.

## Competing Interests

InstaGRAAL is owned by the Institut Pasteur. The entire program and its source code are freely available under a free software license.

## Funding

This research was supported by funding to R.K. and S.M.C. from the European Research Council under the Horizon 2020 Program (ERC grant agreements 260822 and 638240, respectively) and from Agence Nationale pour la Recherche (JPI-EC-AMR STARCS ANR-16-JPEC-0003-05) to R.K. This project has received funding from the European Union’s Horizon 2020 research and innovation programme under the Marie Skłodowska-Curie grant agreement No 764840

## Author’s contributions

LB rewrote and updated the GRAAL program originally designed by HMN, CZ, and RK. MM and AC performed experiments. LB performed the scaffolding. LB and RK analysed the assemblies, with contributions from NG, YLM, AC, KA, LS, JMC, and SMC. LM and RK wrote the manuscript, with contributions from MM, NG, JMC, MC and SMC. LB, MM, SMC and RK conceived the study.

## Acknowledgements

We thank our colleagues from the team, especially Cyril Matthey-Doret, as well as Hugo Darras, Heather Marlow, Francois Spitz, Jitendra Narayan, Jean-François Flot, Jérémy Gauthier, Jean-Michel Drezen, and all Github users and contributors for valuable feedback and comments.

## Supplemental Information

**Table S1:**
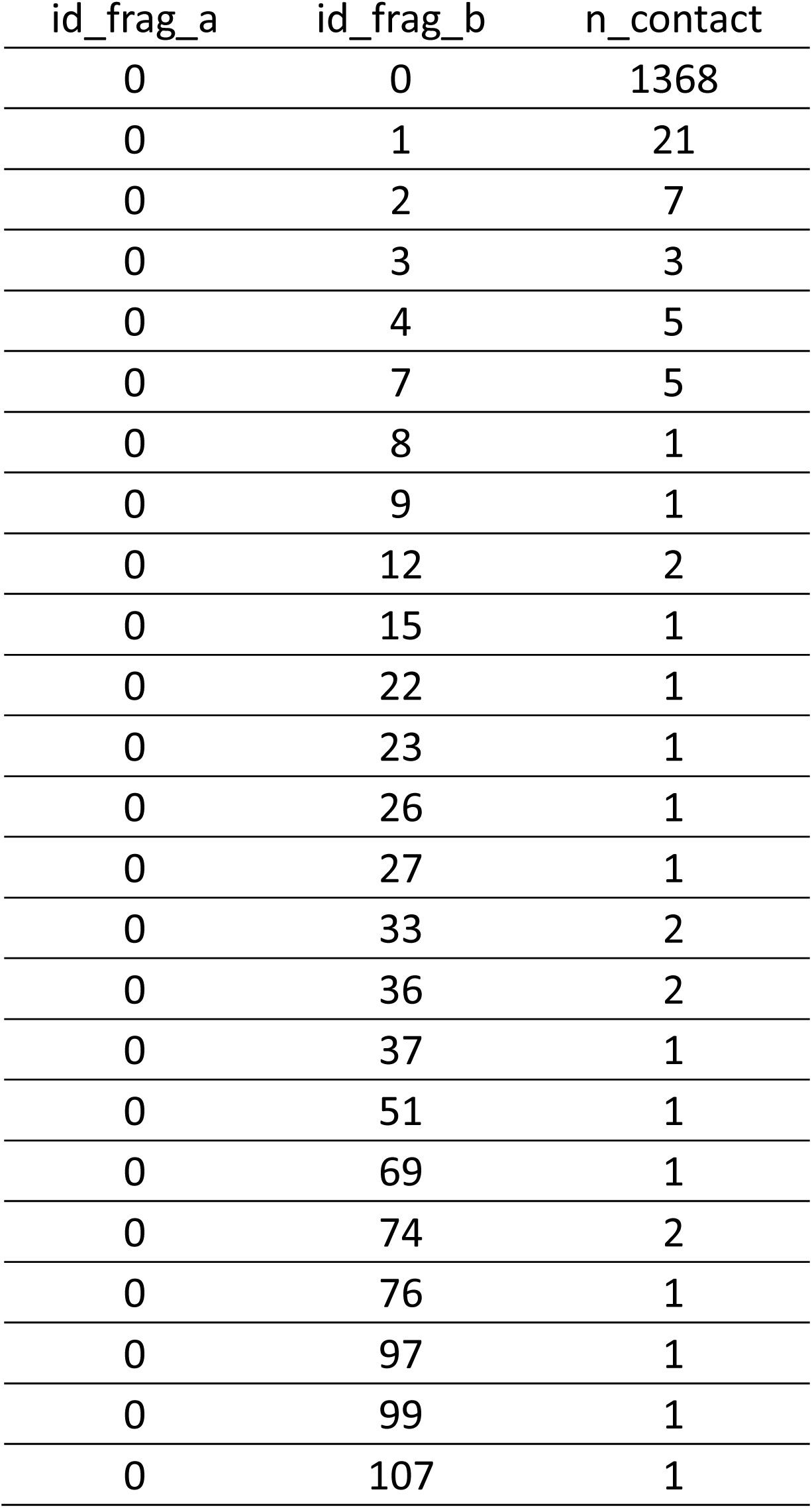
example of a sparse matrix.

**Table S2:** comparison of the integrated sequences between the different assemblies and the reference genome for *Ectocarpus* sp.

|  | Reference<br>genome | Pseudochromosomal<br>assembly | instaGRAAL<br>assembly |
| --- | --- | --- | --- |
| Scaffolds integrated into<br>pseudochromosomes (out of 1561) | 325 | 531 | 793 |
| Percent sequence data Integrated<br>into pseudochromosomes | 70.10 % | 90.50 % | 96.80 % |
| Integrated oriented scaffolds in the<br>(pseudo)chromosomes | 12 % | 49 % | 100 % |
| Number of (pseudo)chromosomes | 34 | 28 | 27 |

**Table S3:**
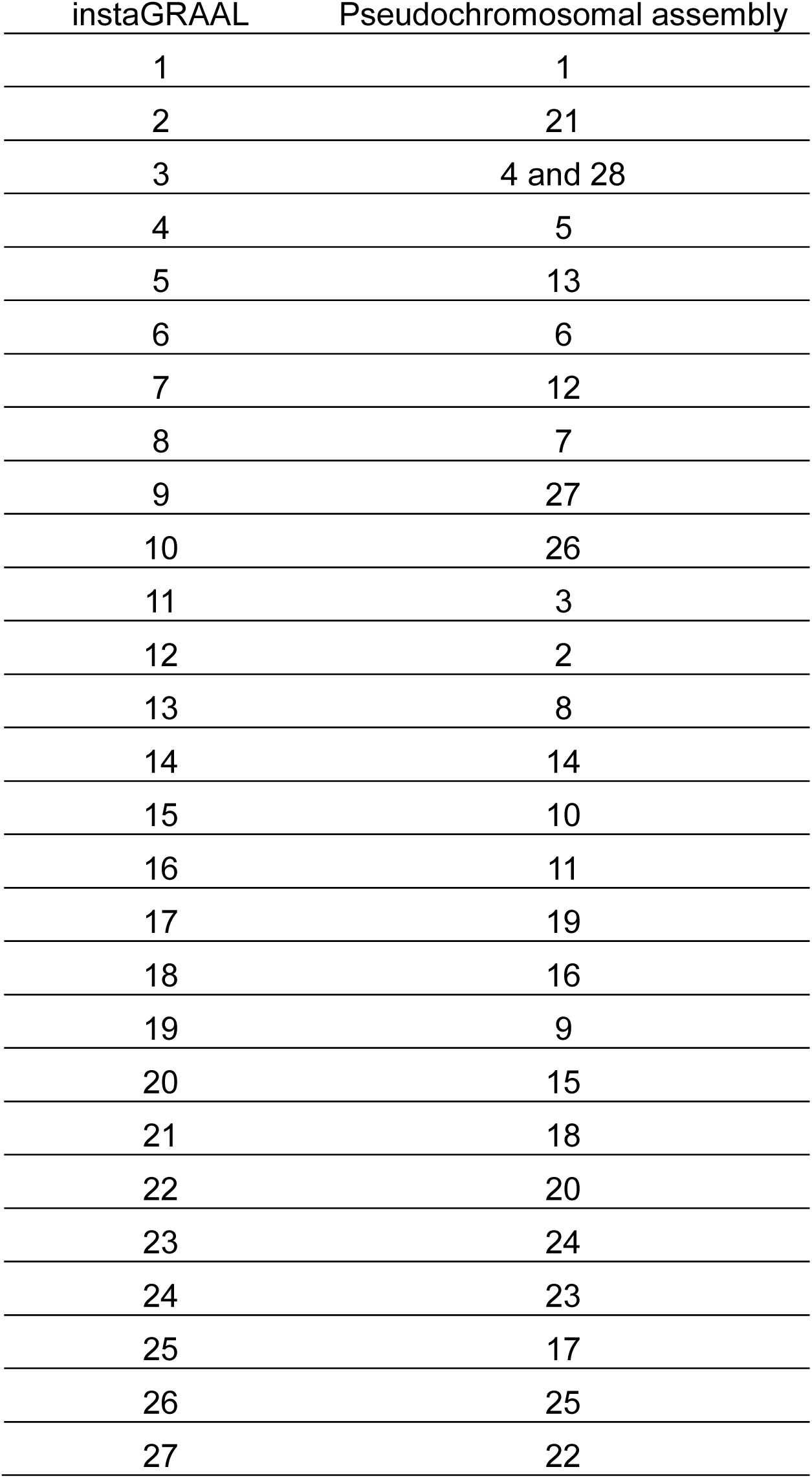
correspondences between instaGRAAL super scaffolds and pseudochromosomes for the *Ectocarpus* sp. genome.

| instaGRAAL | Pseudochromosomal assembly |
| --- | --- |
| 1 | 1 |
| 2 | 21 |
| 3 | 4 and 28 |
| 4 | 5 |
| 5 | 13 |
| 6 | 6 |
| 7 | 12 |
| 8 | 7 |
| 9 | 27 |
| 10 | 26 |
| 11 | 3 |
| 12 | 2 |
| 13 | 8 |
| 14 | 14 |
| 15 | 10 |
| 16 | 11 |
| 17 | 19 |
| 18 | 16 |
| 19 | 9 |
| 20 | 15 |
| 21 | 18 |
| 22 | 20 |
| 23 | 24 |
| 24 | 23 |
| 25 | 17 |
| 26 | 25 |
| 27 | 22 |

**Table S4:**
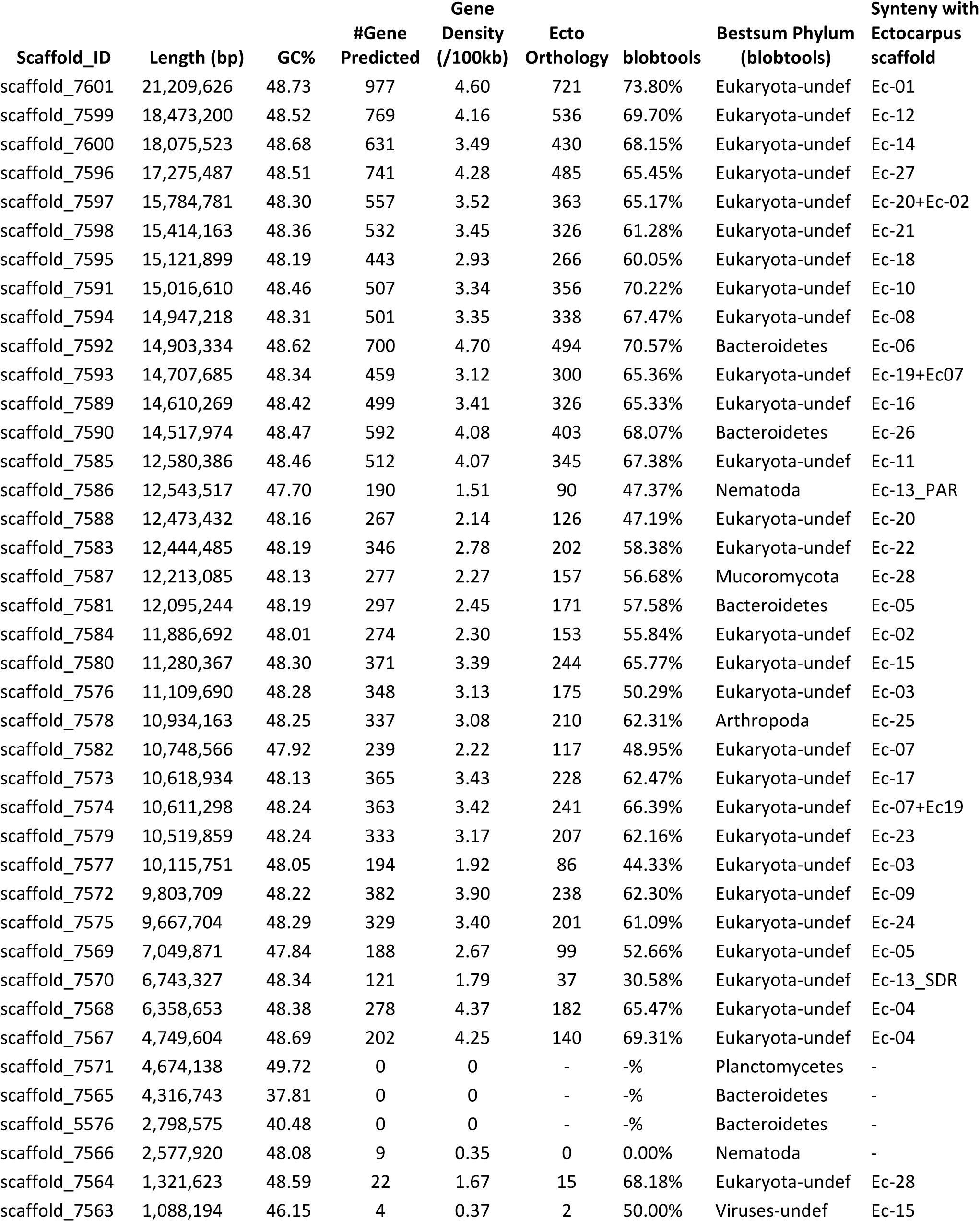
annotation of the 40 *Desmerastia* scaffolds larger than 40Mb.

**Table S5:** metrics of *Desmarestia herbacea* scaffolding using three different programs.

|  | Reference | 3D-DNA | SALSA2 | instaGRAAL |
| --- | --- | --- | --- | --- |
| N50 (bp) | 184,092 | 175,000 | 12,780,148 | 12,444,485 |
| L50 | 697 | 545 | 11 | 17 |
| contig count | 7,743 | 5385 | 4,827 | 4,304 |
| BUSCO completeness | 72.6 | 70.7 | 73.6 | 73 |

**Figure S1:**
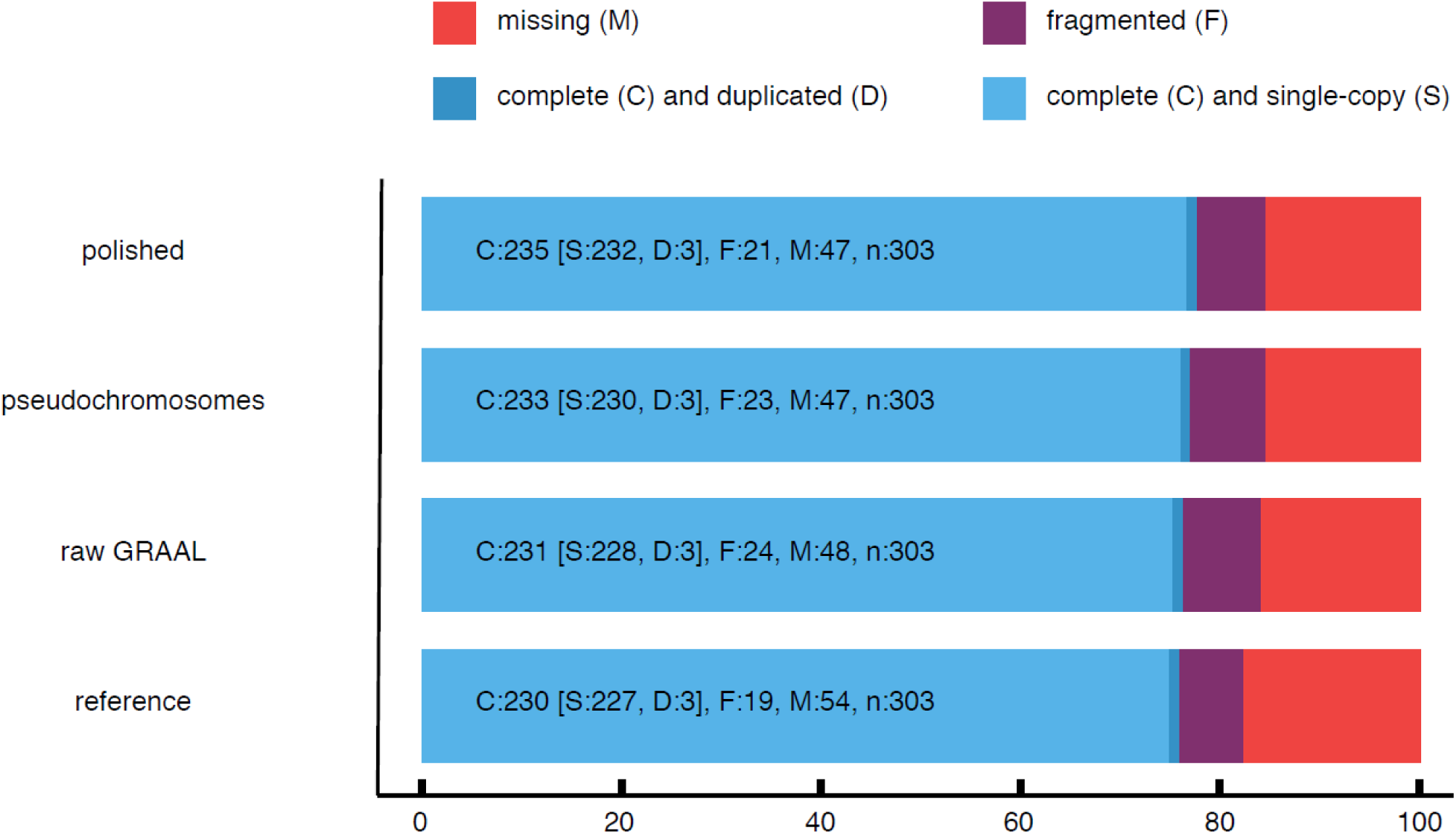
estimates of BUSCO-completeness for the three *Ectocarpus* sp. assemblies and the reference genome.

**Figure S2:**
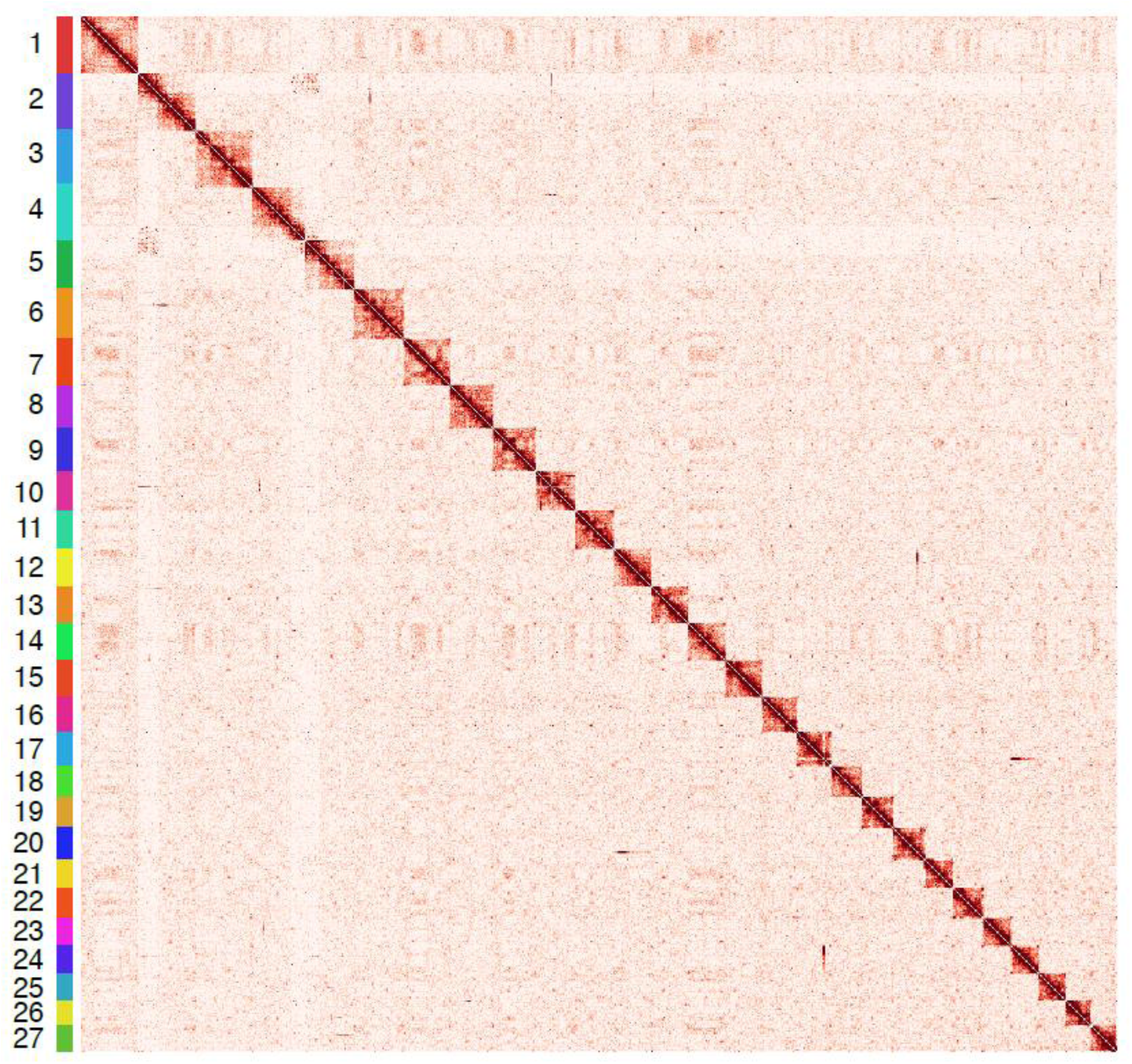
Normalized contact map of the *Ectocarpus* sp. genome scaffolds (presumably corresponding to chromosomes) using instaGRAAL (bin = 200 kb). The colour scale represents the normalized interaction frequencies.

**Figure S3:**
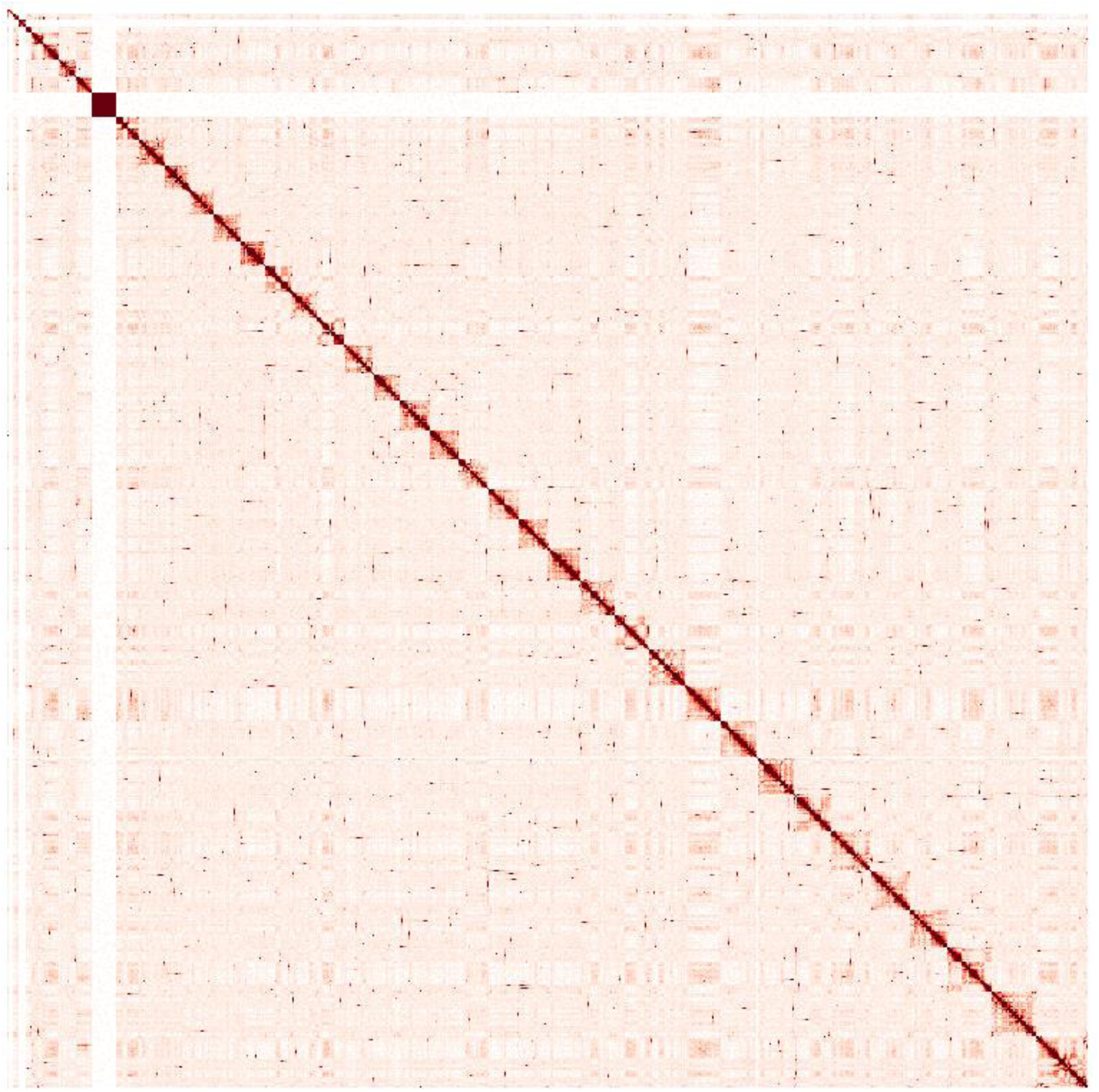
The 40 main scaffolds of *Desmarestia herbacea* after instaGRAAL scaffolding.

**Figure S4:**
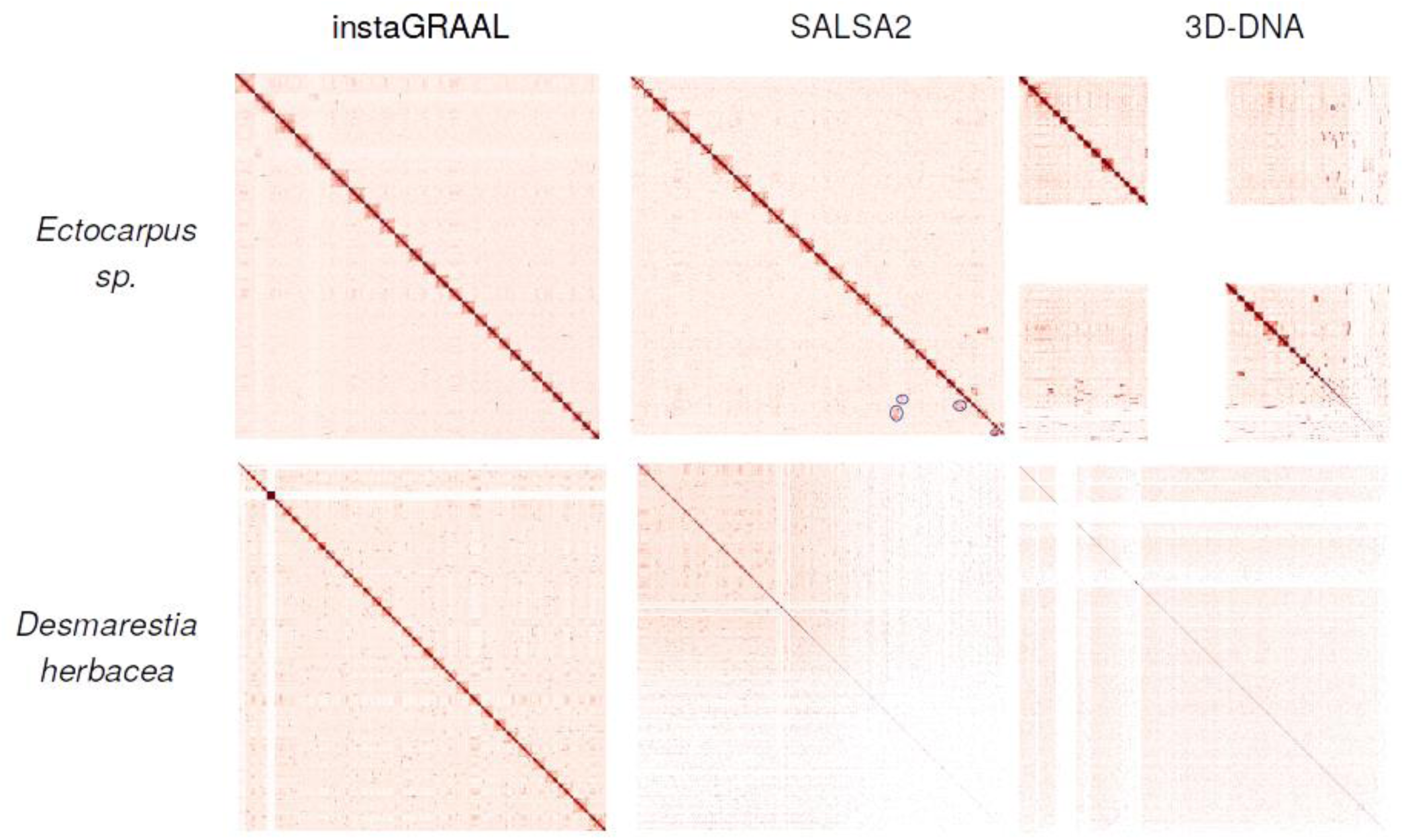
Comparisons of contact maps for three different scaffolders across two species. Gaps represent repeated sequences. Scaffolding differences between the SALSA2 vs. instaGRAAL contact maps on *Ectocarpus sp.* are underlined with circles in the SALSA2 map.

**Figure S5:**
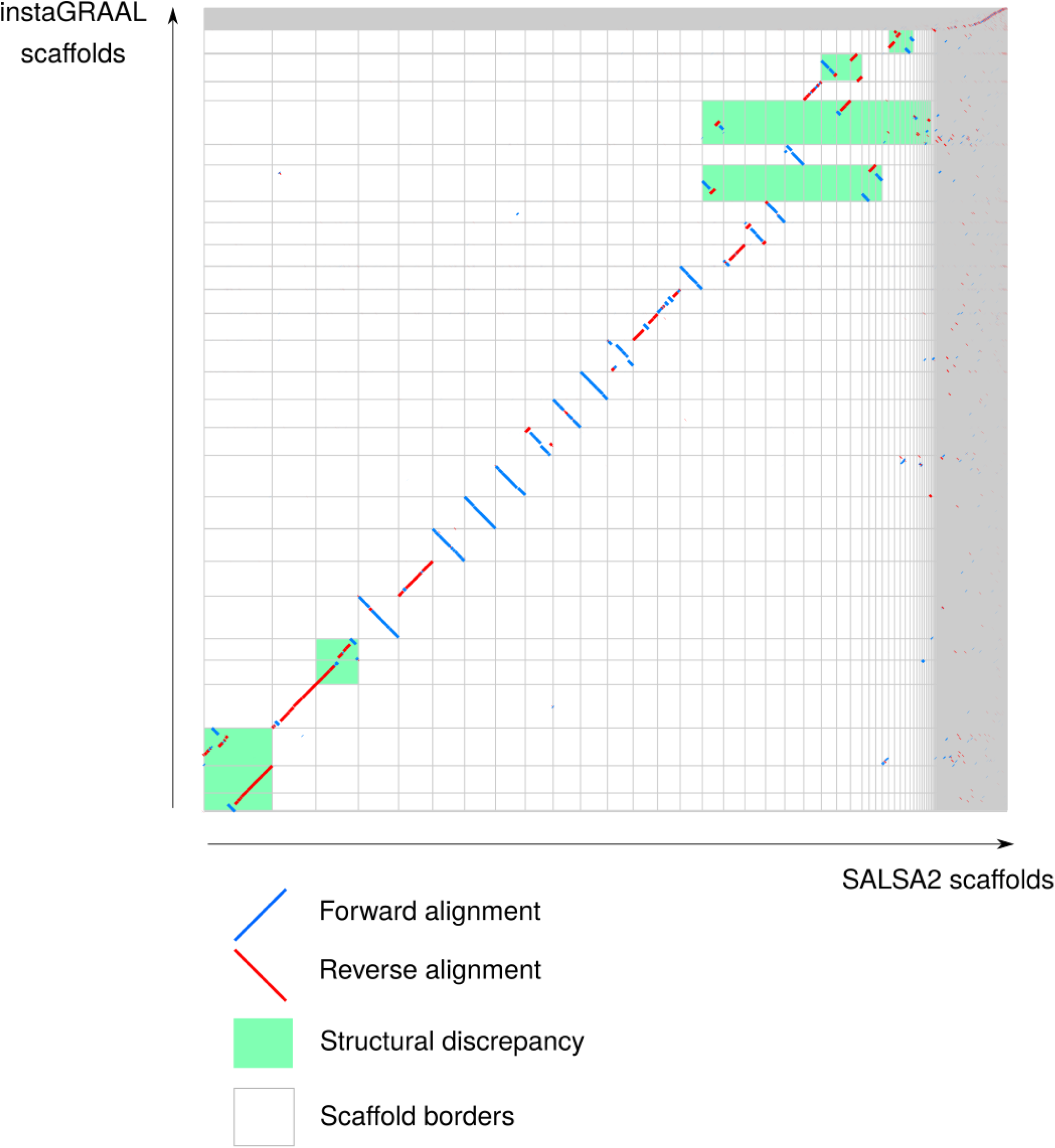
Similarity dotplot of the SALSA2 vs. instaGRAAL 27 scaffolds for *Ectocarpus sp.* Large-scale structural discrepancies have been underlined in green. The contact maps suggest instaGRAAL solutions are more likely.

**Movie S1**: the iterative scaffolding process can be visualized on a movie accessible through the following link. https://github.com/koszullab/ectocarpus_scripts/blob/master/images/matrix_evolution.gif Each frame corresponds to a cycle during which each fragment has been processed once.

## Notes

https://github.com/koszullab/instaGRAAL

https://github.com/koszullab/ectocarpus_scripts

